# Novel Staphylococcal Cassette Chromosome composite island (SCC-CI) with a new subtype of *SCCmec*VI cassette found in ST5 MRSA in France

**DOI:** 10.1101/336396

**Authors:** Olivier Barraud, Frédéric Laurent, Virginie Dyon-Tafani, Céline Dupieux-Chabert, Michèle Bes, Marie-Cécile Ploy, Fabien Garnier, Patricia Martins Simões

**Affiliations:** CHU de Limoges, Laboratoire de Bactériologie-Virologie-Hygiène, Limoges, France; Inserm, U1092, Limoges, France; Université de Limoges, UMR-S1092, Limoges, France; National Reference Center for staphylococci, Hospices Civils de Lyon, Lyon, France; Department of Clinical Microbiology, Northern Hospital Group, Hospices Civils de Lyon, Lyon, France; International Centre for Research in Infectious diseases, INSERM U1111, University of Lyon, Lyon, France

## Abstract

An emergent kanamycin-susceptible ST5-MRSA lineage has been identified in France. Whole genome sequencing revealed a 40 kb SCC composite island with a mosaic structure including 3 SCC elements: a ΨSCC*cop/ars*, a SCC*Lim88A* with a *ccr*C recombinase, and a novel subtype of SCC*mec* type VI (VIb). This mosaic structure suggests a high recombination rate of SCC elements from distinct staphylococci species.

## Main text

Methicillin resistance in *Staphylococcus aureus* (MRSA) is due to the acquisition of an alternative penicillin-binding protein PBP2a which has a low affinity for ß-lactam antibiotics (1). This enzyme, encoded by the *mecA* gene, is horizontally transferred within staphylococci by a large mobile genetic element named the “staphylococcal cassette chromosome *mec”* (SCC*mec*) (2). SCC*mec* elements integrate into the genome of *S. aure*us within th*e o*rf*X/rl*mH ge*ne*, via recombination events mediated by the site-specific recombinases (*ccr* complex) encoded by the SCC*mec* element itself (3). Majority of MRSA clones detected worldwide belongs to a few common clonal complexes (CCs): 1, 5, 8, 22, 30 and 45 (4, 5). The CC5 is one of the CCs clustering the higher number of MRSA clones that have a worldwide distribution. Each of these MRSA clones harbors distinct SCC*mec* elements such as the ones found in the pandemic ST5-MRSA type II (New York/Japan clone), ST5-MRSA type IV (Paediatrics clone), ST5-MRSA type VI (the New Paediatrics clone), ST5-MRSA type I (EMRSA-3) and ST5-MRSA type I (Geraldine clone) (6–9).

In July 2014, a 90 year-old man was admitted at the Limoges University Hospital Center, France, for a sepsis whose origin was a heel bed sore. Blood cultures were positive with Gram-positive cocci in clusters. They were directly tested for the presence of *S. aureus*/MRSA with the Xpert^(r)^ MRSA/SA Blood Culture kit (Cepheid), as recommended by the manufacturer. The presence of *S. aureus* was confirmed by *spa* gene detection, but the presence of a MRSA was not retained as positive, despite a positive *mecA* PCR result, due to the absence of amplification for the *orfX*- SCCmec junction. The day after, antimicrobial susceptibility testing revealed a MRSA profile (strain LIM88). Whole-genome sequencing (WGS) of LIM88 was performed using the Ion Proton^TM^ system (ThermoFisher Scientific), according to the manufacturer‘s instructions. The reads were assembled using MIRA (Mimicking Intelligent Read Assembly). WGS analysis, using both the SRST2 v0.2.0 (10) and RidomsStaphType v2.0 (Ridom GmbH), revealed that LIM88 belonged to the ST5 genetic background and that it had a *spa*-type t777. Search for site-specific insertion sequences (ISS) characteristic of SCC elements (11) revealed the presence of 4 ISS comprising direct repeat (DR) sequences typical of SCC-like cassettes. Thus, LIM88 carries a SCC composite island of 39.6 kb with 3 SCC elements in tandem: a ΨSCC*cop/ars* element (without recombinase genes), a SCC element with a *ccr*C recombinase (called SCC*Lim88A*) and a SCC*mec* element with a type 4 *ccr*AB complex (Figure 1). The nucleotide sequence of this novel SCC composite island was deposited in the NCBI database under the accession number GenBank KX646745.

**Figure 1.**
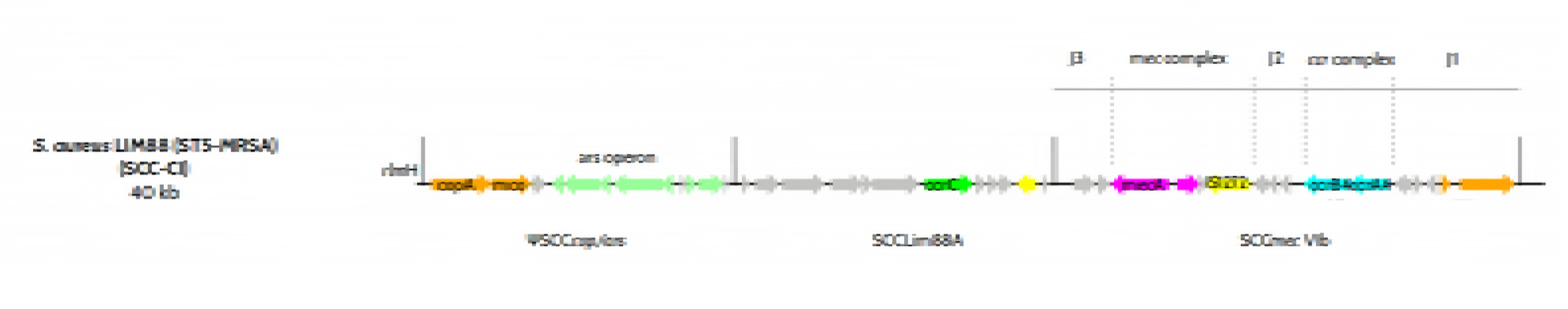
Schematic representation of the SCC composite island ΨSCC_*cop/ars*_-SCC_Lim88A_-SCC*mec* VIb element in ST5 MRSA LIM88. The sequence of the ΨSCC*cop*/*ars*-SCCLim88A-SCC*mec* composite island of strain LIM88 is represented with open reading frames (ORFs) shown as gray arrows indicating the transcription direction. Colored arrows highlight specific genes according to the following criteria: orange = copper resistance related genes (*cop*A and *mco, cop*A*/cad*A); light green = arsenic resistance related genes (*ars* operon); green = staphylococcal cassette recombinase type C gene (*ccr*C); cyan = staphylococcal cassette recombinase type AB genes (*ccr*A4 and *ccr*B4); yellow = transposases (type IS*431* and IS*1272*); magenta = *mec*A complex genes (*mec*A and truncated *mec*R1). The SSC*mec* element architecture with the “non-essential” J1, J2, J3 regions, *mec* complex and *ccr* complex are indicated above the SCC*mec* VIb element of strain LIM88.

Comparison of the 16.6 kb SCC*mec* (ORF25-40, Table S1) element including 16 ORFs found in strain LIM88 with the 23kb type VI SCC*mec* element of the New Paediatric ST5, t777 MRSA clone (strain HDE288) (9) reveals a high nucleotide similarity (>89%) of the “non-essential” region J3, the type B1 *mec* complex (IS*431*- *mec*A-delta*mec*R-IS*1272*), the “non-essential” region J2 and the *ccr*AB (type 4) complex found in both SCC*mec* elements (Figure 2A). Only the J1 regions are distinct (Figure 2A). Interestingly, the J1 region of LIM88 SCC*mec* type VI element shows an overwhelming similarity (more than 90% nucleotide Identity) with a region of a SCC element found in *S. epidermidi*s strain ATCC12228 (Figure 2B), which encompasses the *ccr*A4B4 complex and a *cop*A/*cadA* resistance gene. Based on these data, the SCC*mec* element found in strain LIM88 can be classified as a novel subtype of SCC*mec* type VI element, named SCCmecVIb with the agreement obtained from the International Working Group on the Classification of Staphylococcal Cassette Chromosome Elements (IWG-SCC).

**Figure 2.**
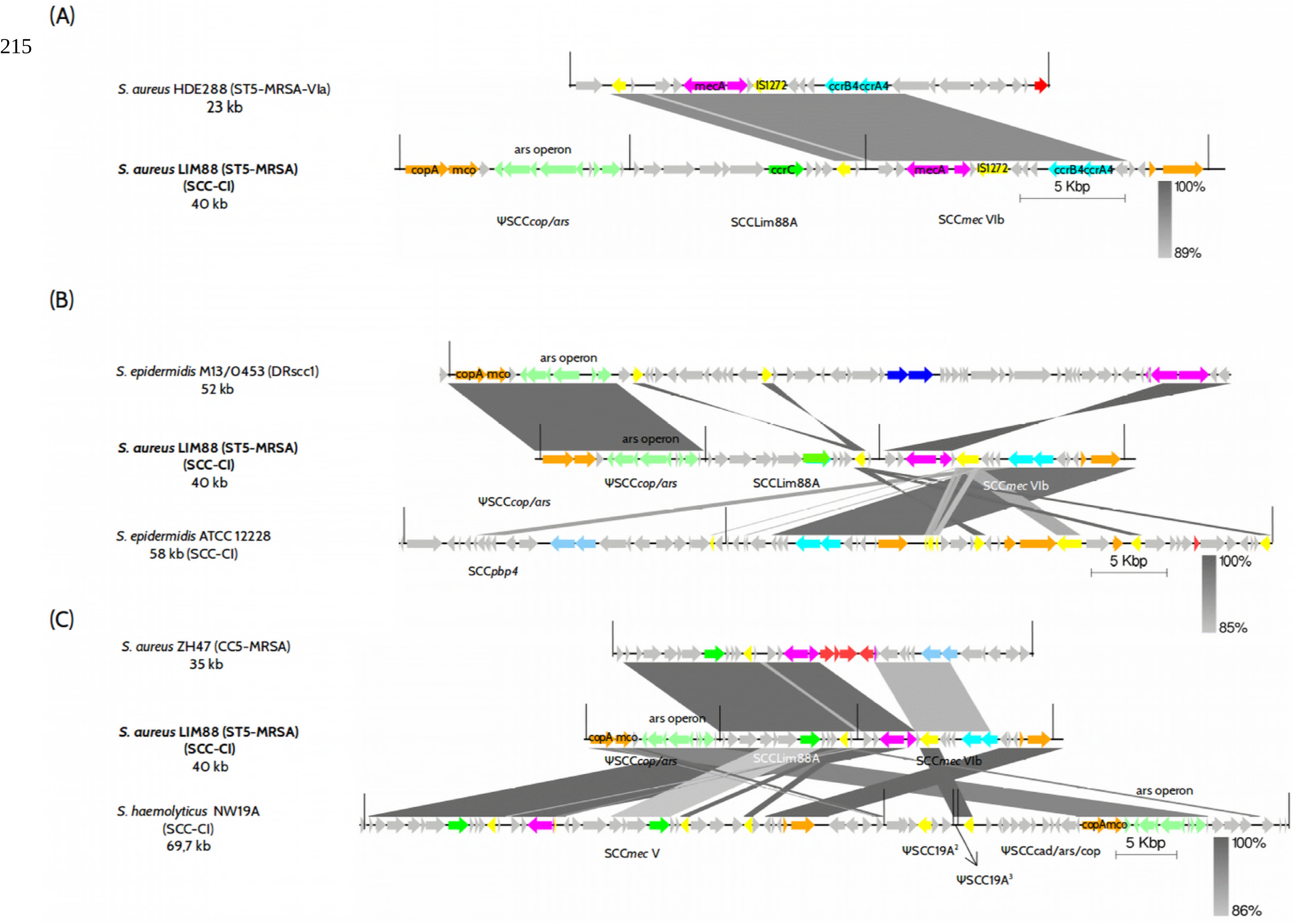
Comparison of the SCC composite island ΨSCC_*cop*/*ars*-_SCC_Lim88A_-SCC*mec* in ST5 MRSA LIM88 with other SCC elements or SCC composite islands (SCC-CI) found in staphylococci. The SCC composite islands or SCC elements are represented with open reading frames (ORFs) shown as gray arrows indicating the transcription direction. Colored arrows highlight specific genes according to the following criteria: orange = copper resistance related genes; light green = arsenic resistance related genes; green = staphylococcal cassette recombinase type C gene (*ccr*C); cyan = staphylococcal cassette recombinase AB (*ccr*A4B4) genes; light blue = staphylococcal cassette recombinase AB (*ccr*A2B2) genes; dark blue = staphylococcal cassette recombinase AB (*ccr*A3B3) genes; yellow = transposases (type IS*431* and IS*1272*); magenta = *mecA* complex genes and red = fusidic acid resistance gene (*fus*) in panel 2A or aminoglycoside resistance transposon Tn*4001* in panel 2B. Homologous gene clusters between SCC elements are indicated by gray diamond areas. The percentage of homology is coded by a gray gradient as indicated in the figure.

Immediately downstream from the SCC*mec* element, a 11.6 kb SCC element (ORF13-24, Table S1) carrying 12 ORFs, among which a *ccr*C recombinase (allele 7) and no known specific virulence or resistance markers, was identified. This element was named SCC_*Lim88A*_. Genomic comparison reveals that this region is highly similar to the *rlm*H proximal J3 region of SCC*mec*_*ZH47*,_ a SCC*mec* element previously described in a CC5 strain isolated from drug-users in Switzerland (12) but, also, to the J3 region found in a composite island of SCC elements (SCC-CI) of the *S. haemolyticus* NW19A (GenBank KM369884) (13), a strain isolated from bovine milk (Figure 2C). This mosaic structure is even more surprising when we look at the SCC*mec* element inserted immediately downstream SCC_*Lim88A*_. The J3 and *mecA* complex itself of the SCC*mec* element are highly similar (90 % nucleotide identity) to the *mec*A complex and its proximal J3 region as found in 3 different *SCCmec* elements: i) the SCC*mec*_*ZH47*_ (SCC*mec* new subtype IV) but without the aminoglycoside resistance transposon Tn*4001* inserted in the *mec*R gene (Figure 2C); ii) the SCC-CI of *S. haemolyticus* NW19A (Figure 2C) and the SCC*mec* type VIa element of strain HDE288 of the ST5-MRSA New Paediatrics clone, as presented in Figure 2A.

Finally, the third SCC element is, in fact, a pseudo-SCC, called ΨSCC_*cop/ars*,_ as it does not carry any recombinase. This 11.1 kb region, located immediately downstream from the *rlm*H gene, harbors 12 ORFS (ORF 1-12, Table S1). Almost all of them encode for heavy metal detoxification functions: two copper related resistance genes (ORF1 and ORF2) and a complete arsenic resistance operon (*ars* operon, ORF4-8, ORF10). As presented in Figures 2B and 2C, this detoxification cluster is present in SCC elements found in other staphylococci species, such as the SCC composite island (SCC-CI) found in the SCC*mec* element of a *S. epidermidis* strain M13/0453 (GenBank MF062491.1) (14) or the SCC-CI of *S. haemolyticus* strain NW19A (GenBank KM369884). The comparison with other simple or composite SCC elements does not help to clarify the origin of this heavy metal resistance cluster as this element is part of canonical SCC elements (carrying within them a cassette chromosomal recombinase) (Figure 2B) but also is part of longer pseudo-SCC elements (SCC elements without the site specific recombinases) (Figure 2C). Nonetheless, the closeness of both functional and truncated IS nearby this heavy metal cluster in almost all SCC elements suggest that they may play a role in the transfer and insertion of this resistance cluster.

No intermediate SCC elements were found in public databases including NCBI or EMBL. The shared similarity with different SCC elements from different staphylococcal species suggests an array of recombination events before the final structure of the SCC-CI, as found in strain LIM88, with no certainty as to the ancestral donor-SCC elements. The presence of this atypical composite island ΨSCC_*cop/ars*_-SCC_*Lim88A*_-SCC*mec*VIb composed by 3 distinct SCC elements, with a non- SCC*mec* cassette inserted at the end of the *rlm*H gene, is likely the reason for the absence of amplification of the *orfX*-SCC*mec* junction when using the Xpert^(r)^ MRSA/SA Blood Culture kit.

We used Kondo‘s PCR (15) primers targeting the different recombinases complexes, *ccrC* and *ccrAB* type 4, of this SCC composite island to perform a preliminary screening of 190 strains of *S. aureus* isolated from blood culture during 2015 in the teaching hospital of Limoges (CHU Limoges), France. The PCR screening suggests that the new ST5-MRSA lineage, carrying the novel SCC-CI ΨSCC_*cop/ars*_*--*SCC_*Lim88A*_- SCC*mec*VIb element, represents more than 15% of blood culture MRSAs isolated in the CHU Limoges. Of note, the positive isolates harbored an atypical antimicrobial phenotypic profile including methicillin and fluoroquinolones resistance and aminoglycoside susceptibility.

In conclusion, we report an original mosaic structure of the staphylococcal chromosome cassette composite island ΨSCC_*cop/ars*_-SCC_*Lim88A*_-SCC*mec*VIb element carried by a ST5 MRSA (strain LIM88) isolated in Limoges, France. Investigations are now ongoing to accurately determine the prevalence and the regional *versus* national distribution of this emerging ST5 lineage in France.

## Acknowledgments

This work was presented at the 2016 ASM Microbe congress in June 2016 (Boston, USA), Poster Saturday-417. We thank the Limoges University BISCEm platform for technical assistance in WGS.

